# Probing the Structure and Functional Properties of the Dropout-induced Correlated Variability in Convolutional Neural Networks

**DOI:** 10.1101/2021.08.19.457035

**Authors:** Xu Pan, Ruben Coen-Cagli, Odelia Schwartz

## Abstract

Computational neuroscience studies have shown that the structure of neural variability to an unchanged stimulus affects the amount of information encoded. Some artificial deep neural networks, e.g. those with Monte Carlo dropout layers, also have variable responses when the input is fixed. However, the structure of the trial-by-trial neural co-variance in neural networks with dropout has not been studied and its role in decoding accuracy is unknown. We studied the above questions in a convolutional neural network model with dropout in both the training and testing phase. We found that trial-by-trial correlation between neurons, i.e. noise correlation, is positive and low-dimensional. Neurons that are close in a feature map have larger noise correlation. These properties are surprisingly similar to the findings in the visual cortex. We further analyzed the alignment of the main axes of the covariance matrix. We found that different images share a common trial-by-trial noise covariance subspace, and they are aligned with the global signal covariance. The above evidence that the noise covariance is aligned with signal covariance suggests that noise covariance in dropout neural networks reduces network accuracy, which we further verified directly with a trial-shuffling procedure commonly used in neuroscience. These findings highlight a previously overlooked aspect of dropout layers that can affect network performance. Such dropout networks could also potentially be a computational model of neural variability.

## 1 Introduction

Convolutional neural networks (CNNs) have been very successful in processing natural images and have been used as visual cortex models (Kriegeskorte, 2015; Yamins and DiCarlo, 2016; Schrimpf et al., 2020). Dropout is a building block of many popular neural network architectures, since it is thought to mitigate overfitting and filter coadaptation problems (Srivastava et al., 2014). In the training phase, a dropout layer randomly sets input values to zeros with a probability (i.e. dropout rate). In the testing phase, the network keeps all input values but scales them down via multiplication by the dropout rate. Some variations of the dropout method also drop values in the testing phase and treat forward computation as sampling the posterior (Maeda, 2014; Kingma et al., 2015; Gal and Ghahramani, 2015, 2016).

Due to the stochasticity of the dropout layers, the activations in the neural networks are no longer deterministic for a given input, but rather are random variables (Fig. 1). The non-deterministic activations have also been widely found in biological neurons: neural responses to an unchanged stimulus fluctuate across trials and through time (Tolhurst et al., 1983). Such neural variability often shows correlated structures among neurons (Zohary et al., 1994; Kohn and Smith, 2005). The correlation coefficient between neural responses to a repeated stimulus among pairs of neurons is known as noise correlation.

**Figure 1:**
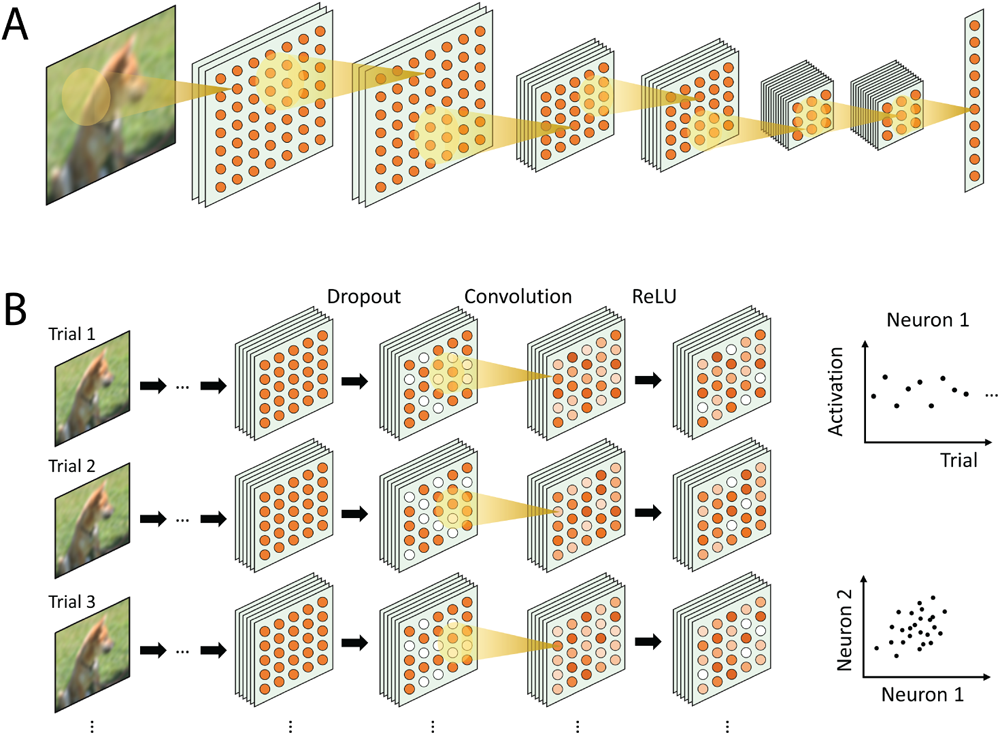
Neural network model and dropout-induced variability. (A) Schematic 6-layer neural network model. It contains widely used building blocks such as 2D convolution, ReLU, Batch Normalization and Max Pooling (see online Methods for more details). (B) Illustration of sampling of the neural network. Dropout is used during both the training and testing phases. We treat each random dropout choice as a trial (a row in the diagram); thus each trial will elicit a unique activation pattern (last column, color indicates level of activation). An example of activations of different trials of one artificial neuron is shown in the top right panel (pseudo data for illustrative purpose). Correlation between artificial neurons can be measured (bottom right panel).

Both empirical and theoretical studies indicate that neural variability can have an important role in neural coding. One prominent view is that noise correlation can significantly impact the amount of information contained in the neural code (Sompolinsky et al., 2001; Averbeck et al., 2006; Shamir and Sompolinsky, 2006; Moreno-Bote et al., 2014; Kanitscheider et al., 2015; Kohn et al., 2016). A decrease in positive noise correlation is generally coupled with an increase in behavior performance (Mitchell et al., 2009; Cohen and Maunsell, 2009; Ni et al., 2018; Gu et al., 2011). Theoretical studies showed that if ”noise” has a component that is aligned with the coding direction of parametric stimuli, the noise correlation is information-limiting (Moreno-Bote et al., 2014), i.e. the amount of information saturates with increasing population size. It is unknown if dropout-induced variability in artificial neural networks is information-limiting and if breaking such structure can improve decoding accuracy, as neuroscience studies would suggest.

The amount of the dropout-induced variance and covariance has been estimated in previous studies (Baldi and Sadowski, 2013; Goel and Chen, 2021). However, an important aspect, namely the structure of the covariance matrix which affects the amount of information encoded, has not been studied. In CNNs, the dimensionality of the response covariance of the whole dataset, but not the within-sample trial-by-trial variability, has been studied and related to the trade-off between efficiency and robustness (Stringer et al., 2019; Nassar et al., 2020; Ghosh et al., 2022; Gerum et al., 2022). On the class level, a study found that the dimension of the manifold spanned by the images within a class decreases in CNNs after training (Cohen et al., 2020). In generative models, low-dimensional geometry and semantic axes have been found in the latent space (Shen and Zhou, 2021; Arvanitidis et al., 2017; Wang and Ponce, 2021). All the above CNN studies that looked into the representation structure and dimensionality of the neural networks are either regarding the whole dataset or classes. The structure and functional role of trial-by-trial covariance regarding each image, which is heavily studied in computational neuroscience, has not been studied on the dropout-induced variability in CNNs. Looking into the structure and dimensionality of the dropout-induced variability can help assess if it reduces information and degrades decoding accuracy, and why it does or does not.

Many dropout variants introduce dependent noise. SpatialDropout drops a whole feature map if one neuron in it is dropped (Tompson et al., 2015). This process presumably can eliminate using gradients from spatially correlated neurons. DropBlock drops contiguous regions from a feature map to efficiently remove semantic information (Ghiasi et al., 2018). Corrdrop drops contiguous regions that have a low correlation with other regions to avoid losing the label information (Zeng et al., 2020). These methods were shown to perform better than the original dropout, but the noise they introduced to the networks is not independent. Another related study is (Dutta et al., 2018), which injected hand-crafted correlated noise into the first layer of a CNN and evaluated the role of noise on the classification of occluded and non-occluded images. We, on the other hand, try to use fewer assumptions and ask if structural neural variance can emerge from only independent and identically distributed (i.i.d) noise. If so, then all the correlation is due to the weight structure but not the external noise structure, as we explain in detail in the Discussion.

Most studies that use CNNs as visual cortex models only focused on the trial-averaged neural response while neglecting response variability across trials (Kriegeskorte et al., 2008; Cadieu et al., 2014; Cadena et al., 2019; Kindel et al., 2019; Kubilius et al., 2019; Deza et al., 2020; Wallis et al., 2017; Mcintosh et al., 2016; Abbasi-Asl et al., 2018). In the brain, trial-by-trial variance can easily be larger than the explainable variance in regression models (Cadena et al., 2019), which arguably is a major difference between the brain and CNNs. There has not been a comparison between CNNs and the brain regarding trial-by-trial variability. Assessing trial-by-trail variability in the dropout CNNs would extend the capability of CNNs as visual cortex models from a new perspective.

In this paper, we study the correlation/covariance structure of the dropout-induced variability in a 6-layer convolutional network trained on the Cifar10 dataset, and test its role in classification performance. First, we find that dropout-induced variability has a similar structure to that found in the brain. Specifically, the noise correlation tends to be positive and low-dimensional, and the main axes of the trial-by-trial covariance matrix are shared between images. Next, we perform two experiments that directly assess the role of correlated dropout variance by breaking the correlation. We find that when we remove the dropout-induced correlation, while maintaining the single-unit variability unchanged, decoding accuracy increases. This increase in accuracy further increases when more neurons are included in the decoder.

Since the correlation structure decreases decoding accuracy, our results could inspire future studies that improve dropout by regulating response covariance. From a neuroscience perspective, the structural similarity we found between the dropout variability and brain variability suggests that CNNs can potentially be used as a model of neural variability.

## 2 Results

### 2.1 Noise correlation is low-dimensional in dropout-induced variability

To assess the sample-wise neural correlation structures induced by dropout layers, we train a 6-layer convolutional neural network with all standard components and training paradigm (See Methods) (Fig. 1A). The channel numbers in the 6 convolutional layers are 64,64,128,128,256, and 256. The center neurons and 3 neurons immediately to the right of the center neuron in each feature map after ReLU are used in the analysis of noise correlation. This population consists of a good combination of neurons at the same spatial location and neurons in the same feature map but at different spatial locations.

The model has dropout layers with a dropout rate of 0.5 throughout the network, but with a modification according to Monte Carlo dropout that drops activations in both the training and testing phase (Maeda, 2014; Kingma et al., 2015; Gal and Ghahramani, 2015, 2016). Due to dropout in the testing phase, the feedforward computations are variable even when an identical stimulus is presented (Fig. 1B). Though not all modern CNN designs have dropout layers after convolution layers and previous studies find that using dropout layers after convolution layers degrades performance (Ghiasi et al., 2018), some state-of-art CNNs still use this design to promote strong regularization (Huang et al., 2017; Tan and Le, 2019; Brock et al., 2021). Since we are interested in dropout-induced variability throughout the network, we choose to put dropout layers after each convolution layer. However, if there is a max pooling layer after the convolution layer, the dropout layer is placed after the max pooling layer to better retain the model average (Wu and Gu, 2015; Wang and Manning, 2013) (See Methods for more details on CNN architecture).

To study the sample-wise neural correlations, we choose to average the following analysis results of the first 100 image samples in the test set. Each image sample is passed to the network 1000 times to get 1000 trials for each image. For each image, we exclude silent neurons in the analysis, i.e. inactive in more than 90% trials. The ratio of the response variance to the response mean, i.e. Fano factor, is slightly smaller than cortical variability, but of a similar order of magnitude (Fig. S1).

The first quantity we look into is the averaged correlation coefficient, i.e. noise correlation, which has been widely studied in neuroscience. In our CNN model, noise correlations are small and positive on average in all layers (Fig. 2B), which is consistent with neurophysiology studies (Cohen and Kohn, 2011). In the brain, noise correlation usually decreases after training (Ni et al., 2018; Gu et al., 2011). To test if the noise correlation is an inherent property of the dropout CNNs or is a result of training, we further look into how training affects noise correlation (Fig. 2A). For early layers (layers 2, 3, and 4), noise correlation gradually increases during training (Fig. 2A). For late layers (layers 5 and 6), noise correlation firstly increases, and then decreases after 15 epochs (Fig. 2A). This result also holds qualitatively when replacing regular dropout layers with DropConnect (Fig. S3).

**Figure 2:**
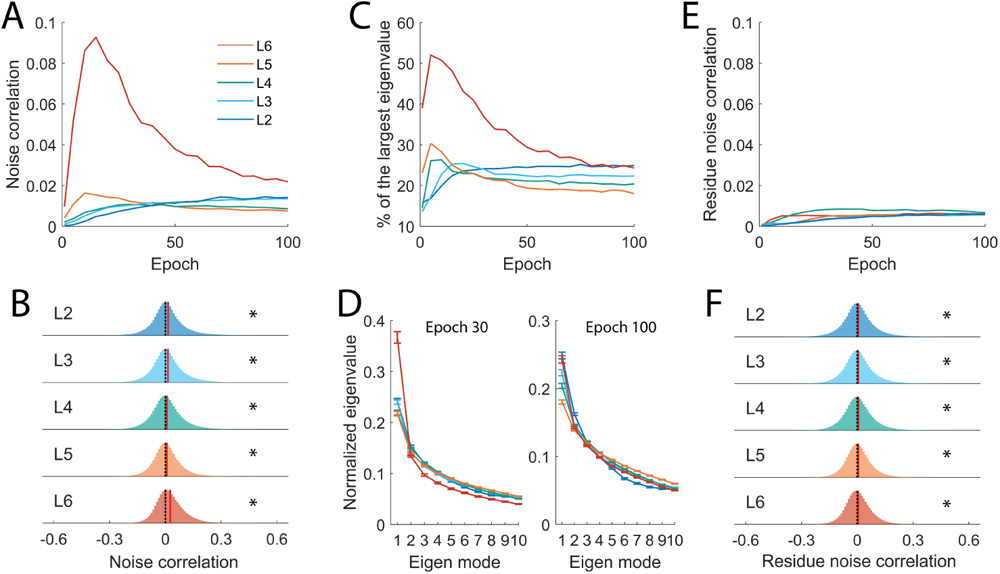
Noise correlation and the dimensionality of the noise correlation in the dropout CNN. (A) Averaged noise correlation in different layers during training. (B) Noise correlation histogram in different layers after training. The dotted line indicates 0; the red line indicates the mean. Asterisks indicate the p-value is smaller than 0.005 in the t-test. Same in later plots. (C) Percentage of the total variance explained by the first eigenmode in different layers during training. (D) The normalized eigenspectrum (divided by the sum of all the eigenvalues) of the loading matrix of the factor analysis of the correlation matrix at epoch 30 and 100. (E) The residue noise correlation computed by subtracting the first eigenmode in (D) from the correlation matrix. (F) The residue noise correlation histogram in different layers after training.

The last layer shows the most drastic change. We think this could be due to a combination of several factors. First, as we will show later, the noise correlation is closely related to filter correlation. We find that the last layer has more filter correlation (Fig S6). This could be a direct reason why the last layer has more noise correlation, though the input correlation structure is also influential (Supplementary section: Source of the noise correlation). Second, though all the network parameters share the same learning rate, the effective learning rate differs between layers. The gradient that backpropagates to the early layers can be smaller than the last layer due to multiplying several small numbers in the chain rule. This could explain why the last layer changes more drastically during the training. The noise correlation dynamic we found echoes a hypothesis of three learning phases and the change of neural variance induced by dropout layers (Baldi and Sadowski, 2013), in which dropout-induced variance starts at a low level, then increases with the weights change, and finally converges to a stable value.

Neurophysiology studies showed that noise variability is low-dimensional (Williamson et al., 2016; Huang et al., 2019; Mendels and Shamir, 2018). The direction of maximal variance in the shared covariance matrix (corresponding to the dominant eigenmode, with the largest eigenvalue) can account for up to 70% of the total shared variance in some cortical areas (Williamson et al., 2016). We study the dimensionality of the dropout variance in the hidden layers of our model. We partition the correlation matrix into independent and shared components by factor analysis (Williamson et al., 2016), and then focus on the shared part of the total covariance. To analyze the dimensionality, we eigen-decompose the shared component of the covariance matrix. The eigenspectrum during and at the end of the training is shown in Figure 2D. A similar eigenvalue spectrum has been found in neurophysiology studies (Figure 1 in (Huang et al., 2019)). During the training, the first eigenmode can account for as large as 50% of the total variance (Fig. 2C), which indicates a low-dimensional structure of the neural correlation.

Eigenvalues do not directly give information on whether one eigenmode contributes to a positive correlation on average across neurons. Intuitively, if an eigenvector x has elements with the same sign, either positive or negative, then this mode contributes to positive correlation (non-diagonal elements in *xx^T^*); whereas if x has equally the same number of positive and negative elements, then this mode does not contribute to positive average correlation. No matter how dominant the first eigenmode is, it is hard to know if it is the main source of the positive noise correlation. Interestingly, we find that the percentage of total variance explained by the first eigenmode changes over training epochs similarly to the noise average correlation (Fig. 2A, C). This gives us a hint it is probably the main source of the noise correlation. We, therefore, hypothesize that the first eigenmode is the main cause of the positive noise correlation. We test this by calculating the residue noise correlation from the residue correlation matrix that is subtracted by the first eigenmode (illustrated in Fig. S2). We find that the change of noise correlation during training is reduced by a significant amount and the value is also reduced (Fig. 2E), though they are still significantly positive at the end of the training (Fig. 2F). This result indicates that there is only one direction that drastically drives the change of noise correlation during training and is the main cause of the positive noise correlation. Replacing regular dropout layers with DropConnect does not change this observation qualitatively (Fig. S3).

### 2.2 Noise correlation depends on the distance and similarity between units

In the brain, neurons that are close to each other have higher noise correlation (Smith and Kohn, 2008). We ask which neuron pairs in the dropout CNN have a larger positive noise correlation. We partition our neural population and measure noise correlation for neuron pairs that only contain neurons at the same location (i.e. between channels, Fig. 3A, B), or in the same feature map but are units apart from each other (i.e. within channels, Fig. 3C, D, E, F, G, H). We find that the distance of neuron pairs within channels is an important factor of noise correlation. Neurons close to each other have higher noise correlation (Fig. 3C, D, E, F, G, H). When neurons are 3-units apart, the noise correlation in layers 2 and 3 exhibits a positive trend but is not significantly positive (Fig. 3G, H). Neuron pairs between channels have significantly positive noise correlations but the effect sizes are small compared to neuron pairs within channels and close to each other (Fig. 3A B).

**Figure 3:**
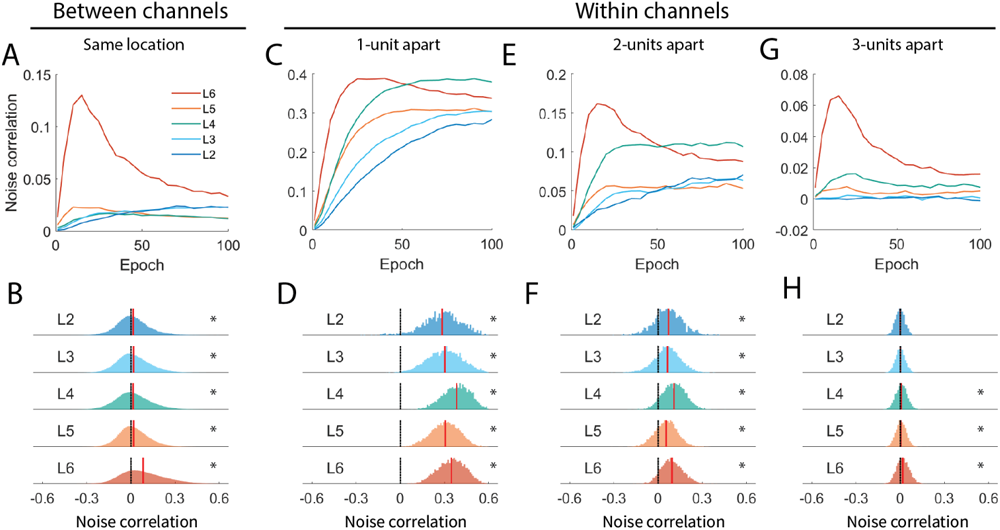
Spatial distance dependent noise correlation in the CNN model. (A) noise correlation of the neuron pairs at the same spatial location but in different feature maps. (B) Noise correlation histogram of the population in (A). (C, E, G) noise correlation of the neuron pairs in the same feature map but at different spatial locations. Neuron pairs are chosen 1-unit apart, 2-units apart, or 3-units apart. We use 1 unit to indicate one neuron in the feature map. (D, F, H) Noise correlation histogram of the population in (C, E, G).

We further test if neuron pairs with higher signal correlation, i.e. correlation coefficient regarding all images, also have higher noise correlation. Neuron pairs in all layers show a significant positive correlation between signal correlation and noise correlation (Fig. S4). A similar relation has been found in neurophysiology studies and named limited-range noise correlation (Zohary et al., 1994; Kohn and Smith, 2005; Gu et al., 2011; Smith and Kohn, 2008). However, the implications of this covariance structure for decoding performance are not trivial, as previous studies have reported diverse effects of limited-range correlations on the encoded information (Kanitscheider et al., 2015; Kohn et al., 2016; Azeredo da Silveira and Rieke, 2021). Therefore, in the following sections, we will directly assess the role of correlated trial-by-trial variability in CNN.

### 2.3 The alignment of the noise covariance and signal covariance

The noise correlation limits the amount of information encoded when it has a component that aligns with the signal direction. This is well studied in cases of parametric stimuli (Moreno-Bote et al., 2014; Rumyantsev et al., 2020a). Note that in those studies the information amount is usually referred to as the Fisher information between neural responses and the parameter of parametric stimuli. In this study, since we focus on the classification task on non-parametric natural images, we refer to the ability to decode the image class from CNN responses as a measure of information. Then, an analogy of the coding direction is the major axis of the centroids of images. Dropout variability that aligns with this major axis could more often intersect between images and cross the decision boundary of the decoder. To measure the alignment of the covariance matrix, we use a metric based on the correlation between the eigenvalues of one covariance matrix and the projection of the other covariance matrix on the first one’s eigenmode (Wang and Ponce, 2021) (See Methods).

We first measure the alignment of the noise covariance between images. The alignment of dropout-induced noise covariance matrices of different images is significant in all the layers after training (Fig. 4A, B). This indicates the local structure of the noise covariance is partially preserved (a perfect alignment is 1 in Fig. 4A) though the images are different (Fig. 4 C). In other words, there is a shared structure of the noise covariance. To define the global neural representation structure, we first calculate the centroids of each image, and then use the covariance of the centroids as an analogy of the signal covariance (Fig. 4F). The alignment of the noise covariance and signal covariance is also significant for all the layers after training (Fig. 4D, E). We further look at how the covariance matrix changes for individual images. We compute the alignment of the noise covariance at different epochs and at the last epoch (Fig. 4I). The alignment becomes high and stable shortly after the beginning of the training (Fig. 4G, H).

**Figure 4:**
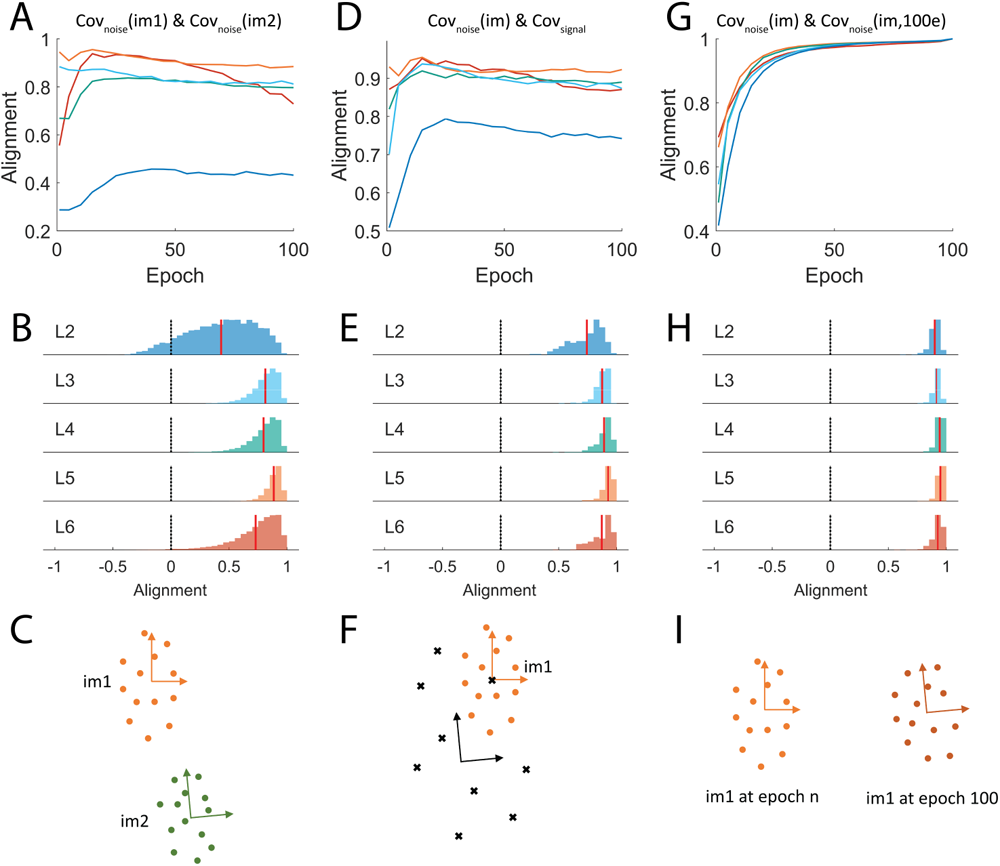
Covariance alignment analysis. (A) Averaged dropout-induced noise covariance alignment between stimuli in different layers during training. (B) Histogram of the alignment in (A) after training. (C) An illustration of the alignment in (A, B) which compares noise covariance axes between different stimuli. (D) Averaged alignment between the signal covariance and noise covariance of different stimuli. (E) Histogram of the alignment in (D) after training. (F) An illustration of the alignment in (D, E) which compares noise covariance axes and the covariance axes of the centroids, i.e. signal covariance. The black cross indicates a centroid of an image calculated by averaging the trials. (G) Averaged noise covariance alignment between epochs and the last epoch of the same stimuli. (H) Histogram of the alignment between the 20th epoch and the last epoch. (I) An illustration of the alignment in (G, H) which compares noise covariance classification task on non-parametric natural images, we refer to the ability to decode the image class from CNN responses as a measure of information. Then, an analogy of the coding direction is the major axis of the centroids of images. Dropout variability that aligns with this major axis could more often intersect between images and cross the decision boundary of the decoder. To measure the alignment of the covariance matrix, we use a metric based on the correlation between the eigenvalues of one covariance matrix and the projection of the other covariance matrix on the first one’s eigenmode (Wang and Ponce, 2021) (See Methods).

### 2.4 The dropout-induced noise correlation reduces decoding accuracy

To directly evaluate the role of the low-dimensional noise correlation and aligned dropout covariance, we conduct two experiments that compare decoder performance of the correlated dropout covariance and uncorrelated dropout covariance. To get the uncorrelated dropout covariance, we run the feedforward computations certain times, and then permute the activations by randomly assigning trial numbers to neurons.

In the first experiment, we stack an SVM classifier after layer 6 (to replace the dense-layer classifier). We control the number of neurons that are recruited in the SVM classifier. To reduce some computations while training multiple models, only center neurons, which presumably contain the most information, are used in this experiment.

The SVM classifiers are trained and tested under two conditions: leaving layer 6 neurons unchanged (correlated), or permuting by assigning artificial neural activations random trial numbers to break the correlations (shuffled). The SVM classifiers trained and tested on correlated activation show lower accuracy and discriminability index (i.e. d prime) than those trained and tested on the permuted activation, and this difference increases as more neurons are used in the classifier (Fig. 5). This indicates noise correlation in the CNN reduces accuracy. This may be related to the information-limiting effects, because noise correlations in our model originate from the feedforward propagation of dropout “noise” (Kanitscheider et al., 2015), although we cannot rule out other factors, given that discriminability for the shuffled data also appears to grow sublinearly. To further show that noise correlation can reduce decoding accuracy in the CNN, instead of training new SVM classifiers on top of the CNN model, we keep the CNN model unchanged. We compare the model accuracy of four different ways of using the dropout layers (Fig. 6): unshuffled trial accuracy, shuffled trial accuracy, Monte Carlo dropout accuracy, and classic use of dropout accuracy (see Methods for details). Shuffled trial accuracy is calculated by trial-shuffling the layer 6 activations to break noise correlation, while the unshuffled trial accuracy keeps the original noise correlation. We use the term ”trial accuracy” to indicate that all the trials of a single image are included in the pool to compute the accuracy.

**Figure 5:**
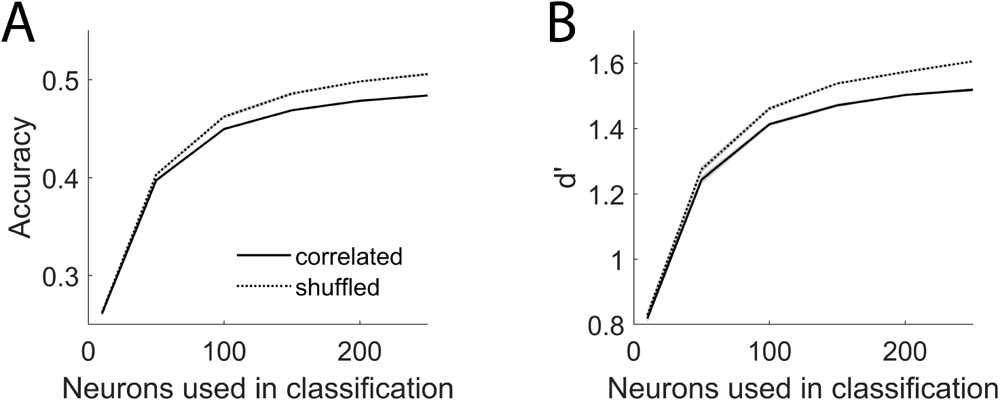
Decoder performance when the layer 6 dropout correlation is eliminated. (A) Accuracy of SVM classifiers trained on different numbers of correlated L6 center neurons (solid line) or permuted to break correlations (shuffled condition; dotted line). When more neurons are recruited, the improvement in the shuffled condition becomes larger. (B) Same as (A) but measuring d prime.

**Figure 6:**
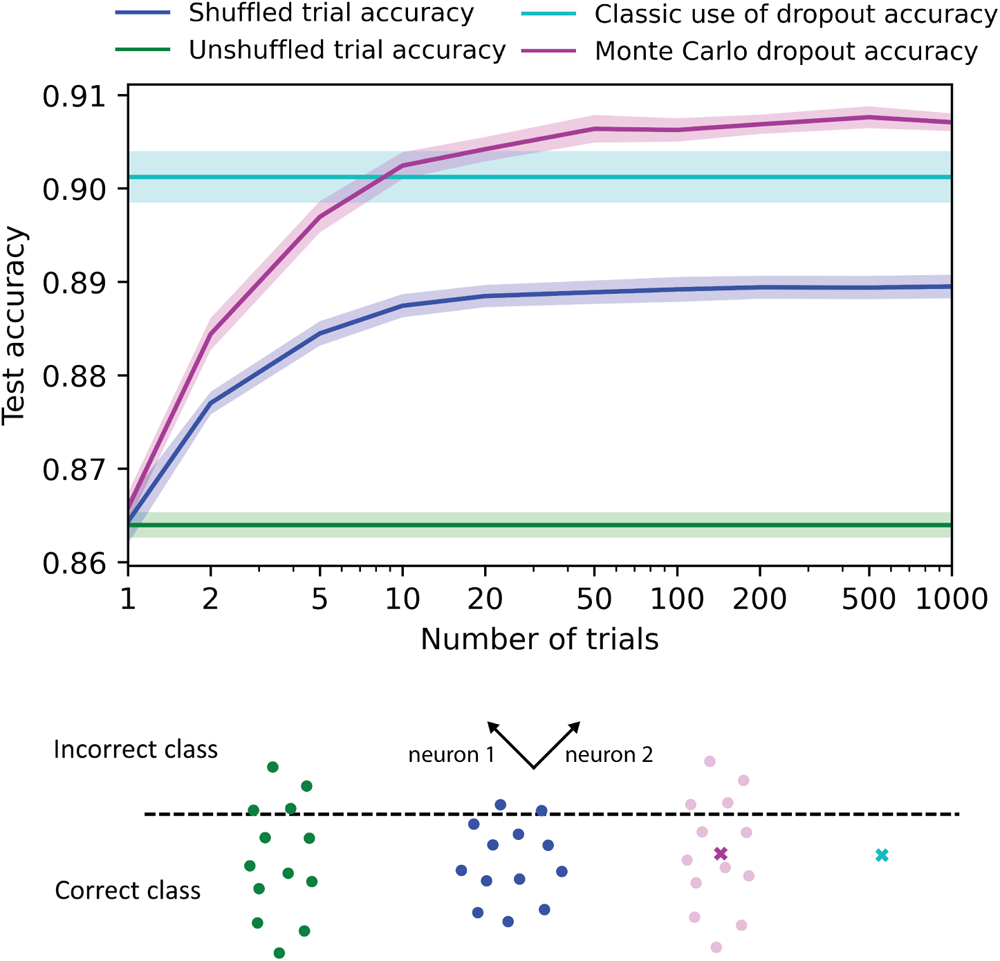
Accuracy difference in four different uses of dropout. Top, network test accuracy of shuffled (dark blue), unshuffled (green), classic use (cyan), and Monte Carlo (magenta) use of dropout layers (See Methods for details). The shaded area indicates the standard error of the mean of 6 models trained on different train-test dataset separations. The unshuffled trial accuracy is the lowest. Breaking the dropout correlation can increase the accuracy (shuffled trial accuracy). However, the two standard uses, classic use of dropout accuracy and Monte Carlo dropout accuracy, of the dropout layers are the best. The shaded area indicates the standard error of the mean of 6 models trained on different train-test dataset separations. An illustration of the four different uses of dropout, for the simplified case of only two neurons and a straight-line decision boundary; in practice, there are more neurons and the decision boundary is a hyperplane. The dashed line indicates a hypothetical decision boundary. The coordinate indicates the hypothetical axes of two neurons. Below the line is the area of correct prediction of this example. From left to right are unshuffled, shuffled, Monte Carlo, and classic uses of dropout. The cross indicates centroid, which is computed in the Monte Carlo case, and is one-step estimated in the classic case. In the unshuffled case, dropout variance has a long-axis towards the decision boundary, which causes more trials to fall out of the decision boundary. In the shuffled case, the dropout variance has a shorter long-axis, thus more trials are on the correct side of the decision boundary. The two standard uses use the centroid to make predictions, thus more likely to be on the correct side of the decision boundary.

The first two ways are the main focus of this study. Comparing them can help assess the role of the structure of the dropout variability. The latter two ways are the current standard uses of the dropout layers. Note that the shuffled trial accuracy and Monte Carlo dropout accuracy depend on the number of trials. For the shuffled trial accuracy, more trials help break the correlation more; for the Monte Carlo dropout accuracy, more trials help estimate the centroids more accurately. We find that the shuffled trial accuracy is higher than the unshuffled trial accuracy, even when only permuting with two trials to break the correlation. This indicates that the correlated structure of the dropout variability indeed reduces decoding accuracy. However, we also find that the two standard uses of the dropout are higher than the shuffled trial accuracy. This suggests that even breaking the correlation could increase trial accuracy. The best use of the dropout layer is still by estimating the centroids and using them to make predictions and interpretations.

Figure 6 shows an intuitive diagram of a simplified case that only has two neurons and a straight-line decision boundary that could explain the accuracy difference between four different uses of dropout layers, though the real case can be more complicated in which there are more neurons and the decision boundary is a hyperplane. The dashed line indicates a hypothetical decision boundary. It is likely to be orthogonal to the main axes of the signal covariance. Since the dropout covariance is aligned with the signal covariance, its main axes are also more likely to be orthogonal to the decision boundary. Breaking the correlation by trial shuffling can shorten the variance on the main axes; thus fewer trials fall on the incorrect side of the decision boundary. The other two standard ways of using dropout layers only use the centroids to make the decision and thus have a higher chance to fall on the correct side.

Does using trial accuracy have any merit over classic dropout and Monte Carlo dropout? Inspired by the diagram in Figure 6, we propose that if there is an adversarial noise that pushes the network output toward the incorrect class, the centroid is pushed to the incorrect side, but in some trials the output remains at the correct class. We conduct an experiment that compares four uses of the dropout layer under adversarial noise (Fig. S5). We find that even with a mild level of noise, the classic use of dropout becomes the worst. With a high level of noise, the shuffled trial accuracy becomes the best.

## 3 Discussion

Inspired by previous computational neuroscience studies, we hypothesize that the structure of the dropout-induced variability in CNNs can affect the network performance. We study the detailed covariability structure in a standard 6-layer CNN model. We find that neuron pairs have significant positive and low-dimensional correlation, and the main axis of the covariance is shared among stimuli and aligned with the signal. The correlation structure we found in the CNN model echoes the findings in the brain. Then, we assess the role of such a correlation structure by comparing network performance of correlated activations and uncorrelated (i.e. shuffled) activations. We find that the shuffled activations indeed outperform the correlated activations, which indicates the structure of the dropout-induced variability degrades decoding performance.

The ”noise” (trial-by-trial variability) in neurophysiology recordings has caught researchers’ interest for a long while. A natural question to ask is if the down-stream brain area can accurately decode the input regardless of the ”noisy” representation. Experimental studies found that such trial-by-trial variability has a positive correlation on average and such a correlation often limits decoder performance (Zohary et al., 1994; Rumyantsev et al., 2020b). Theoretical studies also found that if the correlated neural variance has a component that aligns with the signal, the correlation is information-limiting which means the decoder cannot accurately decode the inputs even with infinite encoding neurons. The above discussion about neural noise is not limited to biological systems. Any stochastic encoder-decoder system can suffer from correlated trial-by-trial variability. Deep neural networks are arguably the most popular artificial encoder-decoder system now, but the structure and the role of trial-by-trial variability have not been studied in detail in such a system. It is crucial to conduct a comprehensive study on the topic, especially given that some studies proposed to include stochasticity during the inference phase, which naturally introduces trial-by-trial variability in the deep neural network.

In electrophysiology studies, cortical noise correlation is reduced under some conditions, such as attention, to increase behavior performance (Mitchell et al., 2009; Cohen and Maunsell, 2009; Ni et al., 2018; Gu et al., 2011). Theoretical studies also suggested that depending on the structure of the correlation matrix, the correlation can benefit the decoding accuracy (Averbeck et al., 2006). In this study, we characterized, for the first time, the structure of the correlation in a dropout CNN model and found it indeed reduces accuracy.

Our study illustrates how noise correlations in CNNs depend on the combination of input noise and network weights. The dropout layer introduces i.i.d. multiplicative Bernoulli noise into the CNN. Since the noise is unstructured, all the trial-by-trial variability structures we observed can only be due to the weight structure. We derive a relationship between output correlation and kernel weight correlation (Supplementary section: Covariance and correlation after dropout layers). If the input random vector is i.i.d., then the output correlation of two neurons is just the filter correlation (i.e. Pearson correlation coefficient between two kernel weights). If the input random vector has a correlation structure, the output correlation is still a similarity measure between two kernel weights but with a linear transformation. We indeed find that in our CNN model, noise correlation is correlated with filter correlation, and filter correlation tends to be positive resulting in positive noise correlation (Fig. S6; see also results on signal correlation in Sec 2.2 and Fig. S4). The low-dimensional noise correlation is usually modeled by low-dimensional global factors (Huang et al., 2019). Interestingly, dropout CNNs without such factors can still show low-dimensional noise correlation. This could be due to the dropout process naturally promoting ”redundant” filters to achieve robustness against dropout noise.

While studying the structure and the role of the trial-by-trial correlation in the CNN model, we also find surprising alignments between dropout-induced variability and neural variability in the brain. Both systems show a positive averaged noise correlation and the correlation is low-dimensional. We further find that the noise correlation is correlated with signal correlation. In both systems, noise correlation can limit decoding performance. This indicates the dropout CNNs could be potentially used as a computational model of neural covariability. The dropout and DropConnect could be thought of as external noise at neuronal and synaptical levels. However, biological covariability results from many mechanisms and sources, including variable inputs (Shadlen and Newsome, 1998), synaptic unreliability (Allen and Stevens, 1994), recurrent network dynamics(Ben-Yishai et al., 1995; Litwin-Kumar and Doiron, 2012; Hennequin et al., 2018), and slow global fluctuations in excitability (Harris and Thiele, 2011; Goris et al., 2014).

Although in this paper we have focused on the information-limiting role of variability, from a different perspective, neural variability can also play a beneficial role that relates to the neural representation of uncertainty (Gal and Ghahramani, 2015, 2016; Buesing et al., 2011; Berkes et al., 2011; Fiser et al., 2010; Echeveste et al., 2020; Festa et al., 2021). According to the neural sampling hypothesis, neural computation is thought to be sampling from the posterior distribution, and a trial is thought to be a sample (Fiser et al., 2010; Orbán et al., 2016). This uncertainty perspective also suggests the importance of studying the dropout variability structure. It is known that the Monte Carlo dropout suffers from miscalibration and cannot estimate uncertainty efficiently(Mitros and Mac Namee, 2019). This is partially because the variational function has a fixed variance. Studies have proposed calibration methods to better capture model uncertainty by adding loss functions that represent miscalibration (Laves et al., 2020; Shamsi et al., 2021). Since our results showed an anisotropic variance, it is reasonable to presume the calibrations of the uncertainty in different directions have different effects. Also, there are studies giving a normative connection between low-level feature uncertainty and neural variability (Orbán et al., 2016; Festa et al., 2021). The theoretical posterior structure derived from such models could be used to guide the regulation of the dropout-induced variability structure for better uncertainty calibration.

## 4 Methods

### Neural network model

The convolutional neural network model is implemented using TensorFlow. We use a 6-layer convolutional neural network with all standard components, such as 2d convolution, max-pooling, batch normalization, and ReLU activation. The model is trained and tested on the Cifar10 image classification dataset (32 x 32 pixels, 10 classes). The channel numbers of 6 convolutional layers are 64,64,128,128,256,256. The kernel weights are initialized using the Xavier uniform method. The kernel size of all convolutional layers is 3-by-3 with “same” padding. The network includes 2-by-2 max-pooling layers after the second, fourth and last convolutional layer. Each convolutional layer is followed by ReLU activation, batch normalization and dropout. A view of the training phase and testing phase behavior of dropout is that computation in the testing phase should capture the average of training phase models. It has been shown that if a dropout layer is placed before a max-pooling layer, multiplying by the dropout rate in the testing phase does not faithfully capture the true model average (Wu and Gu, 2015; Wang and Manning, 2013). Thus, we put dropout layers after the maxpooling layers if there is one after a convolution layer. This design can better retain the model average. Following previous dropout studies, we do not add a dropout layer after the last convolution layer. The networks are trained for 100 epochs by the Adam optimizer with learning rate 0.0001 and L2-regularization coefficients 0.0001. Input images are rescaled so that pixel values range from -1 to 1. A simple data augmentation is applied to the training set (15 degree random rotation, 10% random shift, and random horizontal flip).

The dropout layer is modified so that it drops activations at both the training and testing phase according to Monte Carlo dropout (Gal and Ghahramani, 2016). Only with this modification, the activations in the CNN can have variability to unchanged inputs. The dropout rate is set to 0.5.

### Neural variability analysis

Intermediate layer activations are used for the neural variability analysis. We specifically take the neural activations after the ReLU operation to account for non-negative neural activity. Because we do not use responses immediately after the dropout layer, the variability is not a direct result of multiplicative Bernoulli noise. A better way to think about it is that the variability of each layer’s activation is due to the shared multiplicative Bernoulli noise to its input. A biological view of this can be that, due to various sources of noise, an input neuron failed to respond. If one wants to know the covariance immediately after the dropout layer, it can be derived that the dropout layer scales covariance down by a factor of the square of the dropout rate; the Pearson correlation coefficient after the dropout layer also decreases in an explicit form (see Supplementary section: Covariance and correlation after dropout layers). We also use another dropout variant, i.e. DropConnect, which could be viewed as unreliable synapses, and find the main conclusions still hold (Fig. S3).

Note that the first dropout layer is after the first convolutional layer, so the activations of layer 1 neurons are not variable and not included in the analysis.

In each layer and each feature channel, we include the center neuron and 3 neurons from the center to the right in the following analysis. This population consists of neurons at different spatial locations and also neurons at the same location but tuned to different features. Because some stimuli do not induce a non-zero activation for some neurons, we exclude ”silent” neurons that have no activity in more than 90% of the total trials.

We use the first 100 images in the test set to analyze the variability structure. We are interested in both the local and the global structure of neural response variability. For each image, because of the dropout layers, there is a trial-by-trial variability. We use the term noise correlation and noise covariance to indicate this local, i.e. input-specific, variability. To assess the global structure of neural response variability, we first calculate the centroids of each image from 1000 trials, and then analyze the covariance of the centroids. We use the term signal covariance to indicate this global variability.

### Dimensionality analysis

We use the factor analysis to estimate the dimensionality of neural covariance (Santhanam et al., 2009; Williamson et al., 2016; Huang et al., 2019). Factor analysis partitions variance into a shared component and an independent component, and thus best serves our purpose of characterizing the neural covariance structure. Since the active neuron number varies across stimuli, we choose a constant loading factor number, 10. If the algorithm does not converge on a stimulus (rare occurrence), we exclude this stimulus from the analysis. The shared covariance matrix is constructed from the outer product of the loading factors. The shared covariance matrix is then eigen decomposed, where the proportion of eigenvalues can be interpreted as the percentage of shared covariance explained by that eigenmode.

#### Covariance alignment analysis

We adopt a correlation-based method to measure the axis alignment of two covariance matrices (Wang and Ponce, 2021). The intuition is projecting matrix A to matrix B’s ith eigenmode, then calculating the correlation between the projected values and matrix B’s eigenvalues. If two matrices are only different by multiplying a scale scalar, this alignment measure is 1. On the other hand, if one matrix’s long axis is the other matrix’s short axis, this alignment measure is -1. Note that this naive measure is not commutative regarding two matrices, so we define the alignment by measuring it twice which swaps the two matrices, then takes the mean to be the final alignment measure.

Let the two covariance matrices to be compared be denoted as *C*_1_ *∈ R^n×n^* and *C*_2_ *∈ R^m×m^*. Note that, because we drop silent neurons, the size of the two covariance matrices can be different. The eigendecomposition of the two matrices is *C_i_* = *U^T^* Λ*_i_U_i_*, *i ∈ {*1, 2*}* indicates which covariance matrix, *U_i_* = [**u***_i,_*_1_, **u***_i,_*_2_*, …*], Λ*_i_* = [*λ_i,_*_1_*, λ_i,_*_2_*, …*]. When we project *C*_2_ to *C*_1_’s jth eigenvector, we need to adjust *C*_1_’s eigenvector ac-cording to which neurons are included in *C*_1_ and *C*_2_: **u***^′^*_1_*_,j_ ∈ R^m^, u^′^* ^= *u*^1*,j,l* ^if^ the kth row in *C*_2_ and lth row in *C*_1_ are from a same neuron, *u^′^* = 0 if the neu-ron at kth row in *C*_2_ is silent (absent) in *C*_1_. The projection of *C*_2_ on *C*_1_’s jth eigen-mode is *λ^′^ _′_T* 1*,j C*2u′1*,j*. The directional alignment is then *a_directional_*(*C*_1_*, C*_2_) = *corr*(*λ*_1_*, λ^′^*). To make this measure commutative, we take the average value of the two directions: *a* = (*a_directional_*(*C*_1_*, C*_2_) + *a_directional_*(*C*_2_*, C*_1_))*/*2

#### Analysis of decoding accuracy and information

We use two methods to test if dropout noise correlation indeed reduces decoding accuracy. First, we stack an SVM classifier on top of layer 6 of the CNN model. Center neurons in layer 6 are used as the inputs of the SVM model. Three different subsets of neurons are randomly selected to train SVM models whose accuracy are then averaged and shown in the figure 5. We choose the number of neurons to be 10, 50, 100, 150, 200, and 250, to reveal how performance depends on population size. We either permute layer 6 activations to break trial-by-trial correlation or keep layer 6 activations unchanged. To train the SVM classifier, we randomly select 10000 out of 50000 images from the training set. Each image is repeated 300 times, which is slightly larger than the max neuron number so that a different trial can be used for each neuron in the shuffling procedure. The performance of the SVM classifier is tested on 10000 images of the testing set with the same repetition process. If an SVM model is trained on permuted activation, then it is tested on permuted activation. If an SVM model is trained on correlated activation, then it is tested on correlated activation. Discriminability index (i.e. d-prime) is calculated in the one-versus-all fashion for each class, and then averaged to get a scalar value for each model:

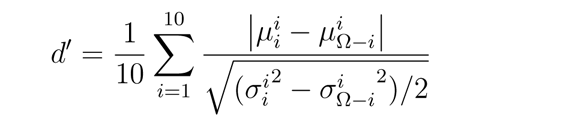

where *µ^j^_i_* is the mean of the predicted probability of class i with true label j; *σ^j^_i_*^2^ is *i i* the variance of the predicted probability of class i with true label j; Ω *− i* is a set of all samples except those with true label i. Note that only center neurons are used in the SVM model. This causes the SVM models to have lower accuracy than the full CNN model which has a dense layer classifier.

Second, we test if only permuting the layer 6 activation and keeping everything else unchanged can boost performance. We compare the model accuracy of using the dropout layers in four different ways (Fig. 6): (1) the trial-wise accuracy that tests with dropout and keeps the neural activation unchanged and dropout variance correlated, i.e. unshuffled trial accuracy; (2) the trial-wise accuracy that tests with dropout but trial-shuffle the layer 6 activations to break dropout variance correlation, i.e. shuffled trial accuracy; (3) the sample-wise accuracy that averages the predictions of different trials of an image, i.e. Monte Carlo dropout accuracy; (4) the sample-wise accuracy that tests without dropout but scales the activation down by half, i.e. classic use of dropout accuracy. We use the term ”trial-wise accuracy” to indicate all the trials of a single image are included in the pool to compute the accuracy; on the other hand, we use the term ”sample-wise accuracy” to indicate there is only one prediction for each image that is included in the pool to compute the accuracy. For the shuffled trial accuracy, all the neurons of layer 6 are permuted by randomly assigning trial numbers to neurons and then passing them to the final dense layer to get predictions. For the first three methods, each image from the testing set is repeated 1, 2, 5, 10, 20, 50, 100, 200, 500, and 1000 times. We do a 6-fold cross-validation by dividing the train and test set in 6 ways and training 6 CNN models.

## 1. Covariance and correlation after dropout layers

The main analysis of this paper is focused on the response variability before the dropout layers. If one is interested in the variability after the dropout layers, it can be derived from the variability structure before the dropout layers. Let *x*_1_ and *x*_2_ be the random variables that represent the responses of two neurons before dropout. Let *m*_1_ and *m*_2_ be the independent multiplicative dropout noise to *x*_1_ and *x*_2_ respectively. *m_i_ ∼ Bernoulli*(*p*), where p is the dropout rate. The responses after the dropout layer are *m*_1_*x*_1_ and *m*_2_*x*_2_. The covariance of the responses after the dropout is:

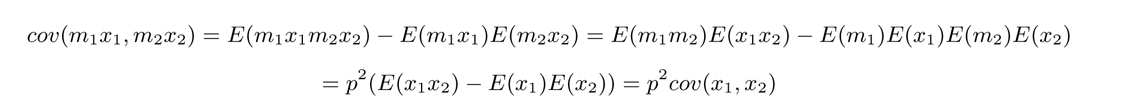

So the dropout layer scales the covariance down by a factor of the squared dropout rate. In our case *cov*(*m*_1_*x*_1_*, m*_2_*x*_2_) = 0.25 *∗ cov*(*x*_1_*, x*_2_) Note that this result does not generalize to the variance since the derivation uses the standard dropout assumption that *m*_1_ and *m*_2_ are independent. The variance after the dropout is:

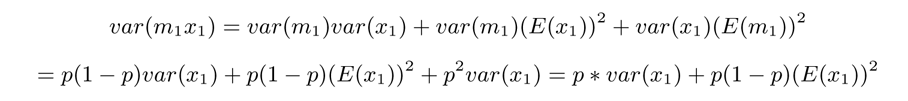

The above equation uses the fact that the variance of a Bernoulli variable is *p*(1 *− p*). In the case of *p* = 0.5, which is a common choice and the choice of our model:

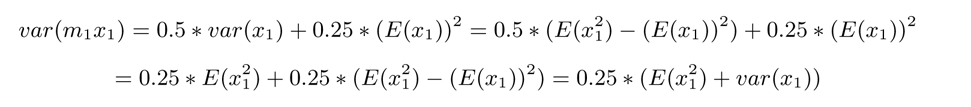

Then the Pearson correlation coefficient after the dropout is:

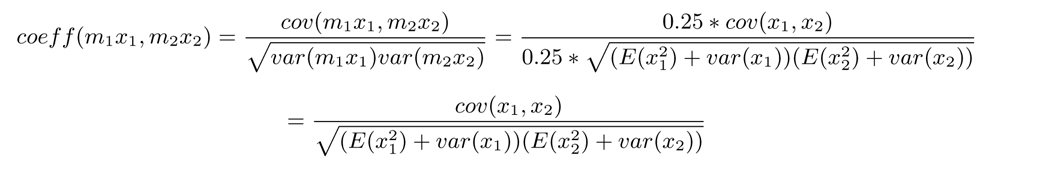

Comparing to,

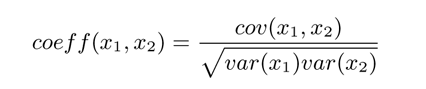

the Pearson correlation coefficient after the dropout decreases. The amount of decrease depends nonlinearly on the mean and variance of the responses.

## 2. Source of the noise correlation

The main finding of our study is that the dropout-induced neural variability has a correlated structure. Given that the dropout noise is independent and does not contain an interesting structure, the observed correlated response noise must be from the kernel (filter) weight structures. Let *x* = *{x_i_}*, *i ∈* [1*…N*] where N is the input dimension, be the input vector of a layer which is a random vector because of the dropout noise, *w*_1_ and *w*_2_ are two kernel weights of two neurons, then the responses of the two neurons are *y*_1_ = *w^T^_1_* and *y*_2_ = *w^T^_2_x*. The covariance of these two neurons is:

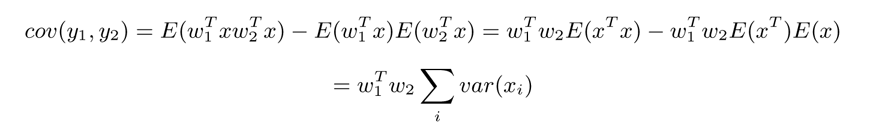

Thus the covariance of these two neurons depends on the product of the two kernel weights. Even when the input random vector *x* does not have any correlation structure, the covariance of the two neurons can be positive when they have a positive inner product; and can be negative when they have a negative inner product.

The Pearson correlation coefficient of the two neurons is:

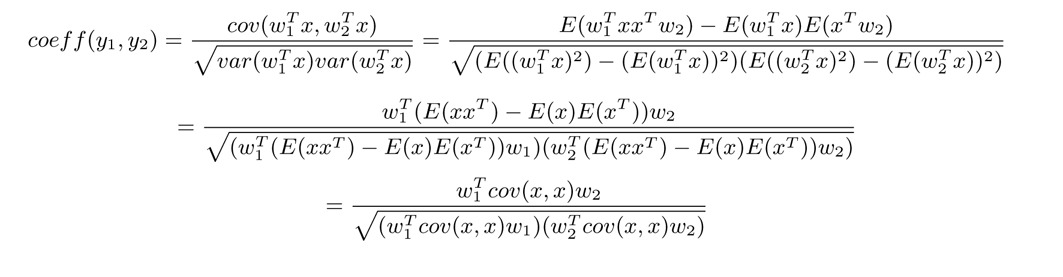

When the covariance matrix of the input is an identity matrix, the Pearson correlation coefficient is:

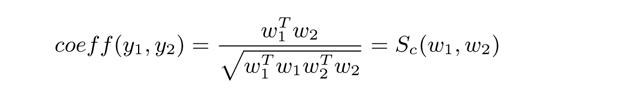

where *S_c_* is the cosine similarity. We call the cosine similarity between two neurons the filter correlation. In the above case, the noise correlation is just the filter correlation.

When the covariance matrix of the input is not an identity matrix, the Pearson correlation coefficient still depends on the similarity between the two kernel weights, but the exact value depends on their cosine similarity after a linear transformation.

We indeed find that the kernel weights are correlated (Fig S4) in the dropout CNN. Thus the correlated kernel weights are the source of the observed noise correlation. We find that the last layer has a higher noise correlation, this could be due to the last layer also having a higher filter correlation, but the alignment between the kernel weights and input covariance matrix could also be a cause.

---

**Fig. S1.**
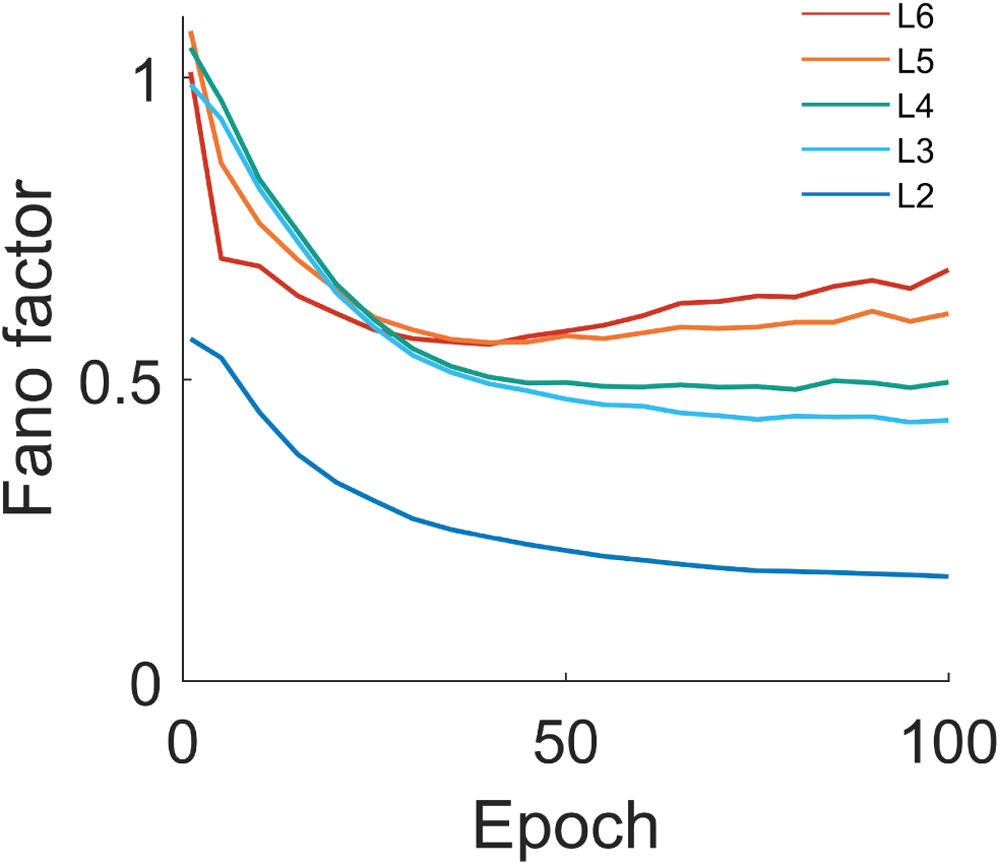
Averaged Fano factors (geometric mean) of different layers during training. The Fano factor is the ratio between response variance and mean, which is known to be close to 1 in cortical neurons. Dropout CNN neurons show a similar magnitude of Fano factor.

**Fig. S2.**
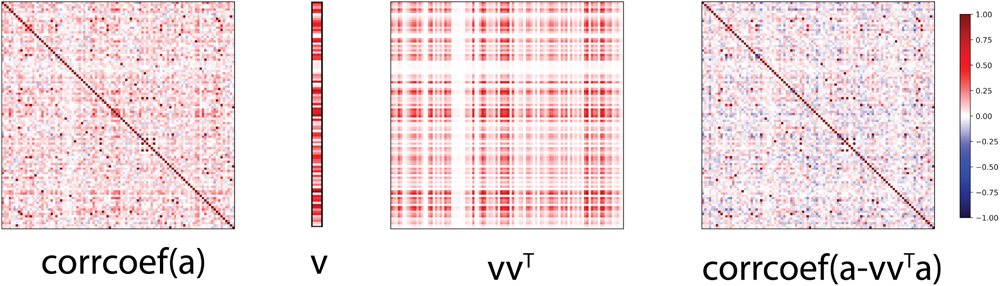
An illustration of subtracting the first eigenmode from the correlation matrix. This is plotted from layer 6 responses elicited by one of the images in the dataset. The First (from left) image is an example of the noise correlation matrix. The second image is the first eigenvector of the shared component of the loading matrix in the factor analysis. Note that, most elements in this eigenvector are positive (red). This indicates it contributes to the positive noise correlation. Its contribution to the correlation matrix is illustrated in the third image. Finally, we project the neural responses onto the null space of the first eigenvector. The reduction of the positive noise correlation can be seen as less red (positive) and more blue (negative) correlations. *a ∈ R^m×n^* denotes the neural activation, where m is the number of neurons, n is the number of trials which in our case is 1000. For a clear display, we only show the first 100 neurons in this layer. The term “corrcoef” denotes the Pearson correlation coefficient matrix with the argument itself.

**Fig. S3.**
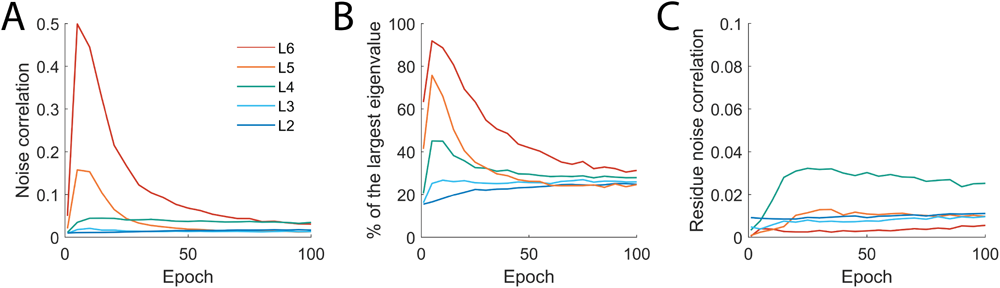
The same analysis as Figure 2 but on a CNN trained with DropConnect. The network architecture and training process is the same as the model used in the main results but without regular dropout layers, and convolutional layers are modified to randomly drop elements in the kernel weights with a probability of 0.5. (A) Averaged noise correlation in different layers during training. (B) Percentage of the total variance explained by the first eigenmode in different layers during training. (C) The residue noise correlation computed by subtracting the first eigenmode from the correlation matrix. The trend and values are qualitatively similar to the dropout CNN model.

**Fig. S4.**
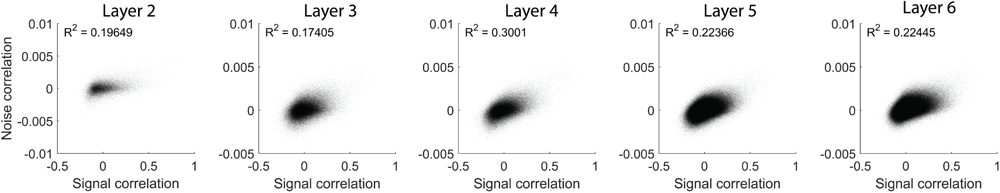
Signal correlation and noise correlation in the CNN neuron pairs. Signal correlation is computed regarding all test images, while noise correlation is computed for each image and then averaged for each neuron pair. In all layers, signal correlation is positively correlated with noise correlation. This type of noise correlation is named limited-range noise correlation in literature.

**Fig. S5.**
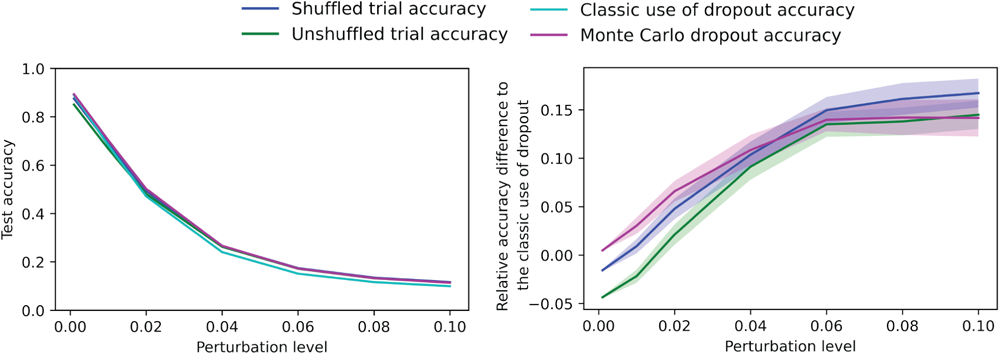
Robustness against adversarial attack. We used the fast gradient sign method (FGSM) to generate an adversarial example for each image. Due to the stochasticity in dropout layers, we computed the gradient regarding the input image for 10 times and used the mean value to generate adversarial examples. Noise levels (perturbation level) 0.001, 0.01, 0.02, 0.04, 0.06, 0.08, and 0.1 were used. The left panel shows the test accuracy of four uses of dropout layers. The right panel shows the relative accuracy difference to the classic use of dropout (subtracting the classic use accuracy, then dividing by the classic use accuracy). A positive number indicates higher accuracy than the classic use of dropout. All other three uses of dropout outperformed the classic use when the noise level is larger than 0.02. Interestingly, the other three uses converge to a similar level of accuracy during perturbation, while the shuffled trial accuracy is slightly higher in extreme noise cases. The shaded area indicates the standard error of the mean of 6 models trained on different train-test dataset separations.

**Fig. S6.**
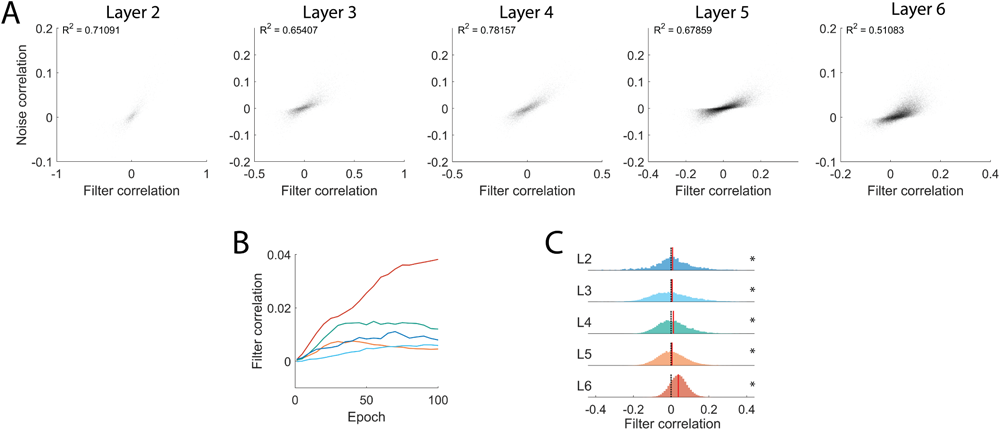
Correlated response noise is due to the correlated filter weights. (A) Noise correlation and filter correlation in the CNN neuron pairs. Filter correlation is defined as correlation between the kernel weights of two neurons. Only center neurons are used in this analysis. We found that noise correlation is correlated with filter correlation. (B) Averaged noise correlation in different layers during training. (C) Filter correlation histogram in different layers after training.

